# A residual memory trace in an accumulator explains serial dependence

**DOI:** 10.1101/2025.08.24.671986

**Authors:** Hiroshi Higashi

## Abstract

Human perception is not a series of isolated snapshots; our recent past continuously shapes what we currently see, a phenomenon known as serial dependence. While convolutional neural networks (CNNs) excel as models of vision, they are typically static and fail to capture such dynamic, history-dependent effects. Here, we introduce SDNet, a model that explains how serial dependence arises from the mechanics of decision-making. SDNet integrates a standard CNN with a recurrent network that functions as a sequential evidence accumulator. We hypothesize that this bias is not a flaw or a feature for optimality, but a natural byproduct of the accumulator retaining a residual memory trace of past decisions. Without being directly fit to behavioral data, SDNet spontaneously reproduces the characteristic patterns of serial dependence from human orientation and numerosity judgment tasks. The model even captures a key feature of human perception: that the bias grows stronger as task difficulty increases. This work provides a concrete, neurally plausible mechanism for serial dependence, challenging theories that frame it as a purely optimal strategy. By showing how a fundamental perceptual bias emerges from an intrinsic property of a dynamic system, SDNet represents a significant advance in building more biologically realistic models of human vision.

## 1 Introduction

Visual perception is not a passive snapshot of the world. It is a dynamic process, profoundly shaped by our recent past. A wealth of evidence shows that our judgement of a stimulus is robustly pulled toward recently seen ones, a phenomenon termed serial dependence [12, 20]. This “attractive” bias is ubiquitous [14, 2, 40], influencing our perception of fundamental features like orientation [15, 25, 43, 45], spatial position [8, 40, 44], color [4, 5, 58], numerosity [14, 18, 21], and shape [39, 38], and extending even to high-level judgement like face identity [36, 55, 56] and expression [30, 37, 60, 17, 41]. Its prevalence suggests serial dependence is not a mere curiosity but fundamental feature of visual processing, yet its underlying mechanism remains a subject a intense debate.

Computational approaches to search for this mechanisms are split between two dominant but incomplete ways. On one hand, traditional cognitive models excel at describing the dynamics of decision-making but are abstract and cannot “see”—they do not operate directly on image pixels [49, 50, 42, 47]. On the other hand, convolutional neural networks (CNNs) have revolutionized our understanding of the brain’s visual pathway and are inherently imagecomputable [33, 34, 61]. However, they are fundamentally static; unlike humans, a standard CNN’s response to an image is deterministic and unaffected by prior stimuli. This leaves a critical gap between biologically-inspired vision models and dynamic human perception.

To bridge this divide, we developed SDNet, an image-computable neural network model that are capable of reproducing sequential effects for image-based inputs. SDNet combines a CNN front-end, functioning as a visual processing system, with a recurrent neural network (RNN) that models the decision process as the accumulation of evidences based on sequential sampling models [10, 11, 23, 29] (Figure 1). Our central hypothesis is that serial dependence is a natural consequence of this accumulation process [44]. Specifically, we propose that the accumulator [9, 28, 59] does not fully reset between decisions, but rather retains a residual trace of the previous choice in its internal memory. This inherited residue, carried in the RNN’s hidden state, then biases the accumulation process for the next stimulus.

**Figure 1.**
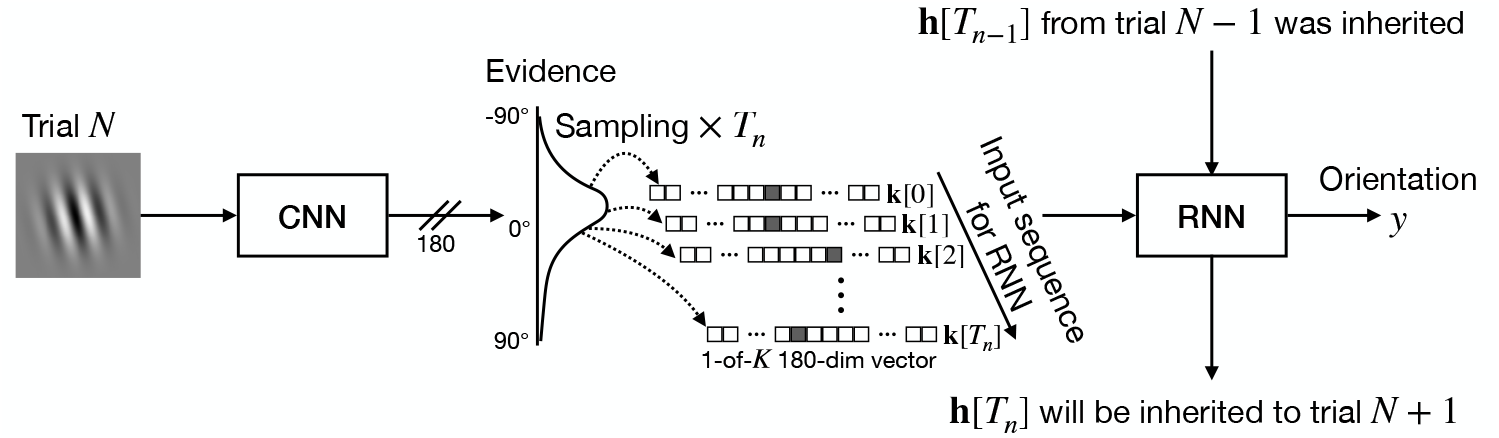
The SDNet model architecture and its mechanism for serial dependence. SDNet’s operation involves three key stages, illustrated here for an orientation adjustment task. (1) Sensory encoding: A CNN processes an input image (a titled grating) and outputs sensory evidence as a probability distribution over possible orientations. (2) Evidence accumulation: An RNN sequentially samples from this evidence to integrate it over time and form a decision. (3) The causal mechanism: Crucially, the RNN’s hidden state is not reset after each decision. Instead, the final state from the previous trial is carried over to initialize the current trial. This inherited memory trace provides a bias that pulls the current decision toward the previous one, thus generating serial dependence.

We tested this hypothesis by presenting SDNet with sequences of images from two classic perceptual tasks: orientation adjustment [20] and numerosity judgement [14, 22]. Without being explicitly trained to do so, SDNet spontaneously reproduced the key signatures of human serial dependence. More impressively, the model also captured the nuanced relationship between task difficulty and the strength of the bias, a hallmark of human behaviour.

This study offers three key contributions. First, we introduce the first imagecomputable model that quantitatively reproduces serial dependence. Second, we provide a neurally plausible mechanism, framing the bias not as an optimized strategy but as an emergent property of a leaky decision accumulator. Finally, SDNet serves as a powerful new tool to forge a more unified understanding of perception, from pixels to dynamic, history-dependent decisions.

## 2 Results

We assessed SDNet’s ability to reproduce core features of human perceptual behaviour by comparing its performance to human participants across two distinct tasks: orientation adjustment and numerosity judgement. We first validated that the model could achieve human-level precision before testing its capacity to spontaneously generate serial dependence.

### 2.1 Orientation adjustment task

We first analyzed an orientation adjustment task (Figure 7) performed by 83 human participants and 83 SDNet instances. In each trial, participants viewed a grating image of random orientation and adjusted a bar to match it. To mimic the varying sensory evidence available to humans, SDNet’s recurrent accumulator processed either 8 or 16 samples from the CNN, analogous to short (0.3 s) and long (0.6 s) stimulus presentation time.

#### SDNet captures human-like performance and its modulation by task difficulty

First, we confirmed that SDNet’s basic performance mirrored that of humans. The average adjustment error was comparable between human (13. ± 5 14.6^°^) and SDNet (15.0 ± 17.8^°^). Critically, both humans and SDNet made larger errors under more challenging stimulus conditions: shorter presentation times and lower spatial frequencies (Figure 2). A repeated-measures analysis of variance (RM ANOVA) confirmed these main effects were significant for both humans (presentation time: *F*_1,82_ = 33.481, *p* < 0.001, 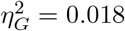 spatial frequency: *F*_1,82_ = 11.023, *p* < 0.001, 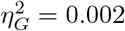) and SDNet (presentation time: *F*_1,82_ = 60.531, *p* < 0.001, 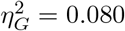 spatial frequency: *F*_1,82_ = 116.379, *p* < 0.001, 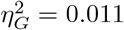). This shows that SDNet’s precision is sensitive to stimulus uncertainty in a manner consistent with human perception.

**Figure 2.**
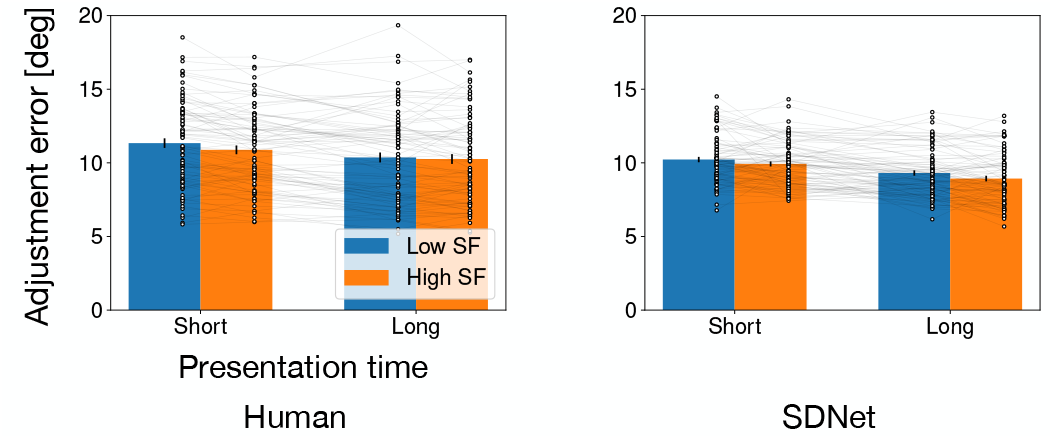
Average adjustment error in the orientation task under different conditions (presentation time and spatial frequency (SF)). Dots represent individual participant/instance means; error bars are standard error of the mean.

#### SDNet spontaneously reproduces the attractive bias of serial dependence

Next, we tested our central hypothesis. Using a derivative of Gaussian (DoG) fit to quantify the bias (see Section 4), we found that both humans and SDNet exhibited a significant attractive pull from the immediately preceding trial (1-back; human: *p* < 0.001, SDNet: *p* < 0.001) and the trial before that (2-back; human: *p* < 0.001, SDNet: *p* = 0.045), as shown in Figure 3. This demonstrates that the model’s core mechanism—the inherited hidden state—is sufficient to generate short-term serial dependence. However, the effect diverged for longer histories: the bias remained for humans at 3-back (*p* = 0.009) but vanished for SDNet (*p* = 0.189), suggesting the model decays rapidly a memory trace.

**Figure 3.**
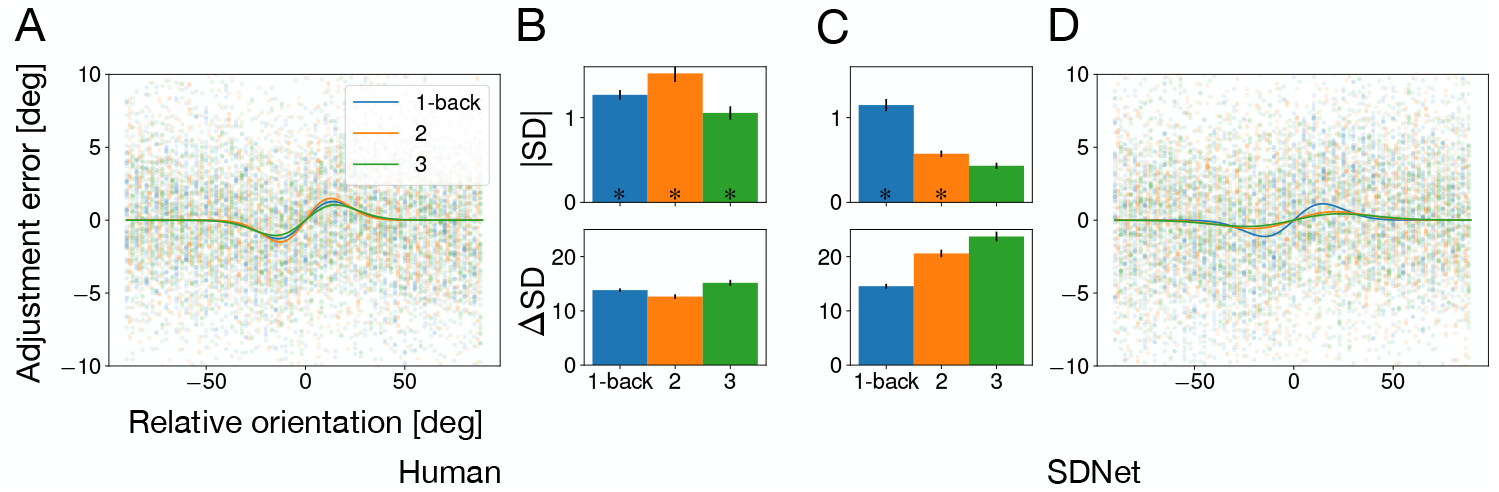
Serial dependence from 1-back, 2-back, and 3-back trials in orientation adjustment. (A) Adjustment errors as a function of relative orientation between the current and preceding stimuli for human participants. Dot represent individual participant means. (B) SD amplitude (|SD|) and peak shift (ΔSD) for human participants. Error bars for |SD| and ΔSD indicate the standard deviation from bootstrap sampling. (C) |SD| and ΔSD for SDNet instances. (D) Adjustment errors for SDNet instances. The asterisks (*∗*) below the bars for |SD| denote significant dependence (*p* < 0.05, bootstrap test).

#### Task difficulty amplifies serial dependence in both humans and SD-Net

A key feature of human serial dependence is its modulation by task difficulty. We found a remarkable correspondence between our model and human data. When the current stimulus was harder to perceive (i.e., had a low spatial frequency), the magnitude of serial dependence significantly increased for both humans (*p* = 0.010) and SDNet (*p* = 0.035), as shown in Figure 4A. This shared pattern supports the principle that when current sensory evidence is unreliable, both the human brain and SDNet rely more heavily on the residual memory of the past. A similar trend was observed for short presentation times (Figure 4B), though it only reached significance for SDNet (*p* = 0.001) and not humans (*p* = 0.682).

**Figure 4.**
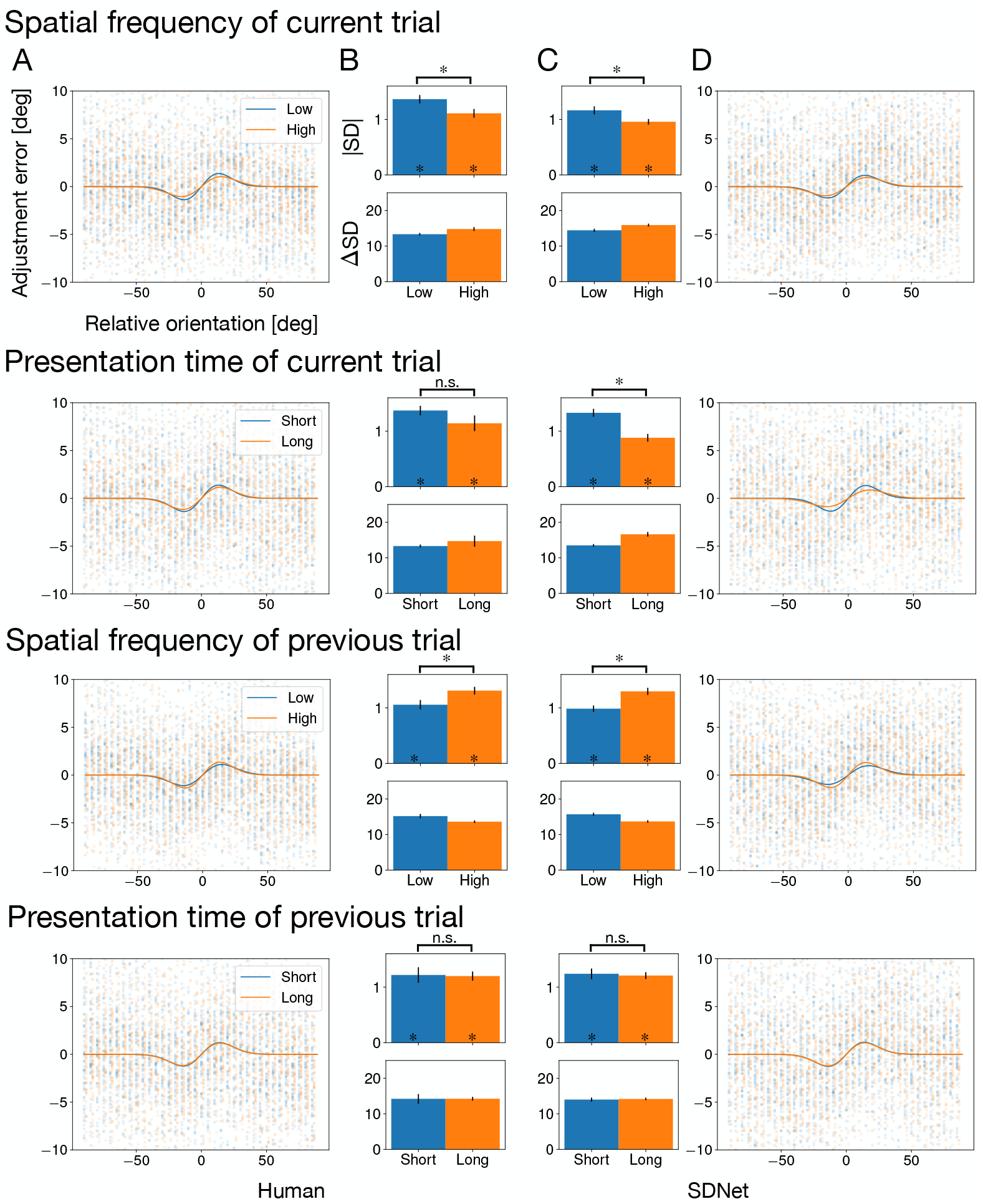
Modulation of serial dependence by stimulus uncertainty. The strength of the 1-back serial dependence bias was conditioned on the properties of the current and previous stimulus. (A, D) Adjustment errors for humans and SDNet as a function of relative orientation. (B, C) SD amplitude |SD| and peak shift ΔSD for humans and SDNet.

#### Previous trial uncertainty similarly attenuates the bias

We also investigated how the reliability of the previous stimulus affected the current bias. Both humans and SDNet showed an identical pattern: when the previous stimulus had low spatial frequency (high uncertainty), it exerted a significantly weaker pull on the current trial (human: *p* = 0.016, SDNet: *p* = 0.001), as shown in Figure 4C. The presentation time of the previous trial had no significant effect for either group (Figure 4D). This demonstrates that both human perception and SDNet dynamically weight past information, discounting it when it originates from a less reliable source.

### 2.2 Numerosity judgement task

To test the generality of our model, we compared it against a public dataset from a numerosity judgement task [22]. In this task, 32 participants judged whether a “probe” image had more dots than a “reference” image, after being primed by an “inducer” image. We likewise trained 32 SDNet instances on the same task.

#### SDNet matches human precision but not all task-specific biases

SD-Net’s overall task precision was indistinguishable from that of humans (*Z*_31_ = 158.000, *p* = 0.238; two-sided Wilcoxon signed-rank test with Bonferroni correction; Figure 5A). However, SDNet did not capture a specific recency bias observed in humans (*Z*_31_ = 60.000, *p* < 0.001; Figure 5B), who tended to respond “more” when the probe and reference were physically identical, a nuance our general-purpose accumulator architecture does not account for. Although this bias led to a slight difference in their psychometric functions (Figure 5C), the points of subjective equality (PSEs) derived from these functions were not significantly different between human and SDNet (*p* = 0.053; bootstrap test (see Section 4.2.3); Figure 5D).

**Figure 5.**
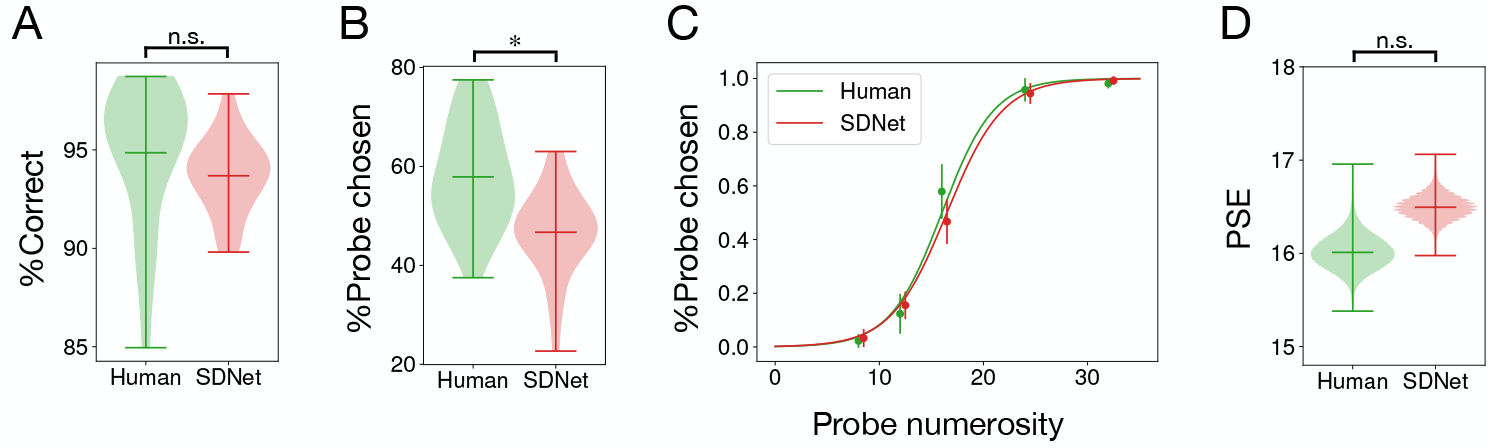
Behavioural performance in numerosity judgement task. (A) Percentage of correct responses for the judgement task. The three horizontal bars represents maximum, average, minimum of PSEs, respectively. (B) Percentage of responses indicating “probe stimulus had more dots” when the reference and probe stimuli had equal numerosity (16 dots). (C) Psychometric functions, plotting the proportion of “probe has more dots” responses as a function of probe numerosity. Dots represent the grand average, and error bars show the standard deviation across participants/instances. (D) PSEs.

**Figure 6.**
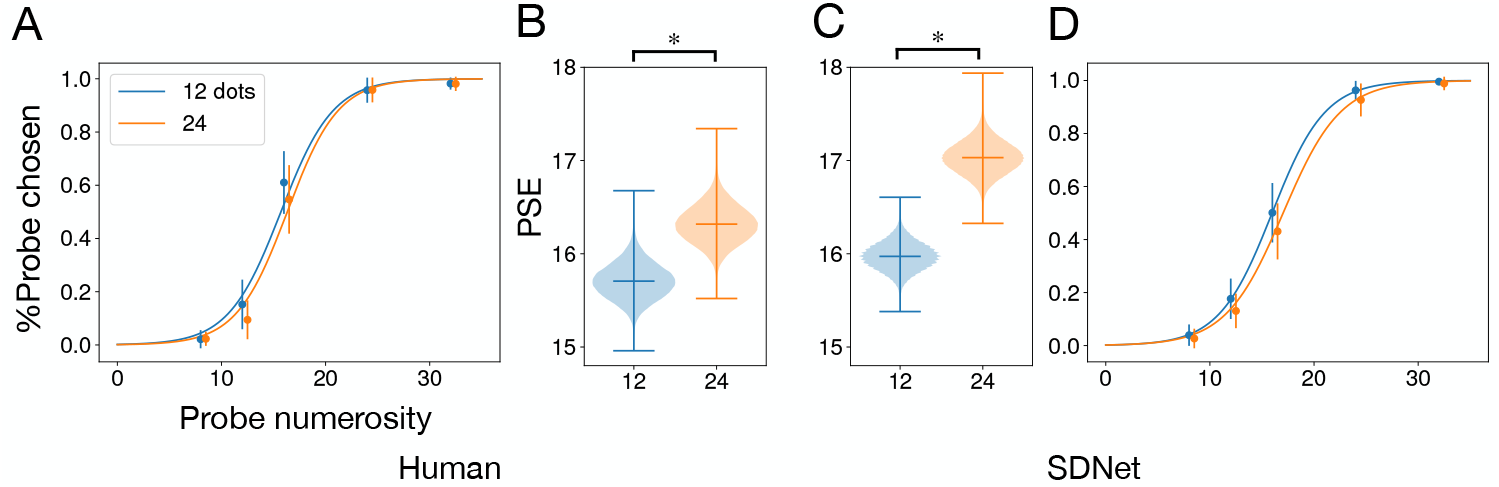
Serial dependence modulated by inducer numerosity in numerosity judgement task. (A, D) Psychometric functions for humans and SDNet, split by low (12 dots) versus high (24 dots) inducer numerosity. (B, C) PSEs for human and SDNet.

**Figure 7.**
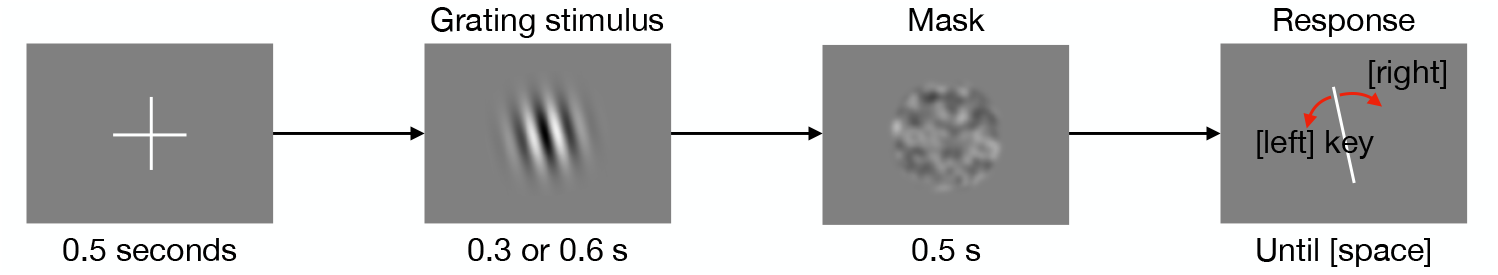
Trial structure of the orientation adjustment task. Participants first viewed a central fixation cross, followed by a brief presentation of sinusoidal grating. After a mask, a response bar appeared, which participants rotated to match the orientation of the grating they had just seen.

#### SDNet robustly reproduces serial dependence

Despite the minor difference in bias, SDNet robustly reproduced the core serial dependence effect. For both humans and SDNet, the judgement of the reference stimulus was significantly biased by the numerosity of the preceding inducer stimulus. A lownumerosity inducer caused the subsequent reference to be perceived as having fewer dots (shifting the psychometric curve left), while a high-numerosity inducer did the opposite (human: *p* = 0.001; SDNet: *p* < 0.001; Figure 6). This successful replication in a different task and stimulus domain strongly supports our central claim that serial dependence is an emergent property of evidence accumulation with a leaky memory trace.

## 3 Discussion

Neural networks are powerful tools for modeling vision, but they have largely overlooked the brain’s dynamic nature, where past experiences continuously influence current decisions. In this study, we addressed this gap by developing SDNet, an image-computable model that spontaneously reproduces serial dependence, a hallmark of human perception. By testing it on orientation and numerosity tasks, we demonstrated that SDNet not only matches human-level performance but also captures the core attractive bias of serial dependence and, critically, how its strength is modulated by task difficulty.

### A mechanistic alternative to optimality

Our work offers a new perspective on the very nature of serial dependence. Prevailing computational theories often frame this bias as an optimal strategy derived from Bayesian inference or predictive coding, designed to enhance perceptual stability and efficiency in a noisy world [18, 16, 42]. SDNet challenges this view. We propose that serial dependence is not a dedicated feature for optimization, but rather an emergent property—a natural side effect of how the brain makes decisions.

Our model’s architecture is built on the well-established principle of sequential evidence accumulation [51, 23, 48, 13, 19]. We hypothesized that the neural accumulators responsible for this process are not perfectly reset after each choice. Instead, a residual memory trace of the previous decision, carried in the accumulator’s internal state, persists and biases the next one [44, 4]. Our results validate this hypothesis, shown this simple mechanism is sufficient to generate human-like serial dependence.

This reframes a key question: instead of asking why the brain evolved an optimal strategy for serial dependence, we can ask why its decision accumulators are not fully reset. The answer may lie in biological constraints, such as the metabolic cost of a hard reset, or functional benefits, like faster processing by starting from a non-zero baseline.

### Bridging models and neurophysiology

SDNet also provides a concrete framework for investigating the neural underpinnings of serial dependence. Neurophysiological studies have found correlates of past stimuli in both early visual areas like V1 [54, 53] and higher-level frontopariental regions [1, 7, 57]. This has fueled debate about where in the processing hierarchy the bias originates.

Our model, with its CNN-to-RNN structure, maps onto this hierarchy. While the CNN “sees” the stimulus (like V1), the bias originates in the RNN “accumulator” (like higher-level cortex). This aligns with findings that decision-related accumulation processes are observed in parietal and frontal cortices [32, 31, 3, 27] and that these regions are causally involved in serial dependence. The influence of these higher areas could then feedback to early visual cortex. SDNet thus provides a powerful tool to test such hypotheses by correlating its internal layer activations with neural recordings from different brain regions [61].

### Limitations and future directions

While SDNet captures core phenomena, its simplicity also highlights clear avenues for future work.

SDNet’s bias decays rapidly, failing to reproduce the weaker but significant dependence observed from 3-back trials in humans. The longer-term dependence likely suggests the involvement of other cognitive systems [52] not included in SDNet, such as working memory [6] or episodic memory. This points to a critical avenue for future research: investigating the interplay of multiple memory systems in human perception [25].

Our model only accounts for attractive biases. However, under certain conditions, perception is repulsed from a prior stimulus [24, 26, 2, 47]. These repulsive effects are often considered a signature of distinct neural mechanisms and represent an important phenomenon for future models to explain.

Serial dependence in human is strongly modulated by the spatial location of stimuli. The current version of SDNet is not spatially organized. A natural next step is to develop an architecture with multiple, spatially-tuned accumulators to investigate these critical spatial dynamics.

### Conclusion

We introduced SDNet, an image-computable neural network that explains serial dependence as an emergent property of evidence accumulation with a leaky memory. This work provides a neurally plausible, mechanistic alternative to optimality-based theories and represents a critical step toward building more comprehensive, dynamic models of human vision that bridge the gap from pixels to perception and decision.

## 4 Methods

This study investigated serial dependence by comparing the performance of human participants with that of a novel neural network, SDNet, across two perceptual tasks. All analyses were conducted in Python, and the model was implemented in PyTorch.

### 4.1 Orientation adjustment task

To validate SDNet’s ability to reproduce human serial dependence, we collected human behavioural data through an online experiment.

#### 4.1.1 Human behavioural experiment

##### Participants

One hundred participants were recruited via Prolific (www.prolific.com). All provided informed consent before the experiment. The study protocol was approved by the Committee for Human Research at the Graduate School of Engineering, The University of Osaka, and adhered to the Declaration of Helsinki. Participants received a base compensation of £3 for completing the experiment, with an additional performance-based bonus ranging from £0 to £1 (average: £0.5) based on their accuracy in orientation adjustment.

##### Stimulus and procedure

Each trial consisted of a sequence: fixation, stimulus presentation, mask, and adjustment phase (Figure 7). Stimuli were sinusoidal gratings using the `GratingStim` module in PsychoPy [46]. For each trial, the grating’s spatial frequency (`sf`) was randomly selected from {2, 3, 4} Hz, orientation (`ori`) from {0, 1, 2, …, 179} degrees, and phase (`phase`) from {0, 0.1, 0.2, …, 0.9} (in steps of 0.1). Stimuli were presented at the center of the browser window, masked by a Gaussian function (`mask: gauss`), and scaled to half the window size. Stimulus presentation duration was either 0.3 s or 0.6 s. A mask image was presented for 0.5 seconds immediately after the stimulus to prevent visual aftereffects.

Participants were instructed to adjust a keyboard-controllable bar to match the orientation of the previously presented grating stimulus. The bar’s initial orientation was randomized. Participants used the left and right cursor keys for counter-clockwise and clockwise rotations, respectively. There was no time limit for the adjustment, and participants finalized their response by pressing the space bar. Each participant completed 200 trials, separated into 4 blocks of 50 trials each, with a mandatory break of at least 30 s between blocks.. The average experiment duration was 27 minutes.

##### Data screening

Initial participant screening excluded data from participants whose average absolute orientation error exceeded 39.2^°^ (calculated as 1.5 × (interquartile range) + (first quantile)). This criterion led to the exclusion of 17 participants (17%). Subsequently, trial-level screening excluded individual trials with an absolute orientation error above 37.5^°^ (1.5 × (interquartile range)+ (first quantile)). This excluded 488 samples (approximately 2.9%). The final dataset for analysis comprised 16,112 samples from 83 participants.

#### 4.1.2 SDNet

##### Architecture

SDNet consists of two primary modules (Figure 1): a CNN for visual feature extraction and an RNN for sequential evidence accumulation and decision-making. The CNN is an AlexNet [35] that takes a 128 × 128 pixel image and outputs a 180-dimensional probability vector representing sensory evidence for each orientation. The RNN is a gated recurrent unit (GRU) with 16 hidden cells. At each time step *t*, it receives a one-hot vector ***k***[*t*] (sampled from the CNN’s evidence) and updates its hidden state ***h***[*t* − 1] to produce ***h***[*t*]. The final hidden state is passed through a dense layer to produce the model’s perceptual report.

##### Training

Training was a two-stage process. First, the CNN module was trained on a large dataset of generated grating images to predict the correct orientation. The dataset of 5,400 grating images (128 × 128 pixels) was generated using PsychoPy’s `GratingStim` module, with parameters matching those used in the human experiment (spatial frequency (`sf`): 2, 3, 4, orientation (`ori`): 0, 1, 2, …, 179, phase (`phase`): 0, 0.1, 0.2, …, 0.9). These images were split into training (3,240 images), validation (1,080 images), and test (1,080 images) sets with a 6:2:2 proportion. The CNN was trained to minimize the cross-entropy loss between one-of-*K* coded ground-truth orientation *y*_*n*_ and its predicted probability distribution ***e***_*n*_. Early stopping was employed: training was terminated when the average absolute error of the CNN module on the validation dataset reached below the human-averaged orientation error (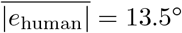).

Following CNN training, the RNN module was trained with the CNN module’s weights frozen. For each training sample, the CNN’s output evidence ***e***_*n*_ was repeatedly sampled (16 times) to generate a sequence of 180-dimensional one-hot vectors ***k***[*t*], *t* = 0, 1, … 15. The RNN module was trained to minimize the cross-entropy loss between the true orientation *y*_*n*_ and the RNN’s final output evidence ***ϵ***_*n*_. Crucially, to model serial dependence, the initial hidden state ***h***_*n*_[0] for the current image ***x***_*n*_ was set to the final hidden state ***h***_*m*_[*T*] of a randomly chosen previous image ***x***_*m*_ from the training set. Early stopping for the RNN training was similarity applied based on reaching the human-averaged orientation error on the validation dataset.

##### Generating behavioural responses

To simulate human behavioural data, SDNet processed sequences of images. For a given trial *n* with input image ***x***_*n*_ and a previous trial *n −* 1 with image ***x***_*n−*1_, the CNN module first computed evidence ***e***_*n*_ for ***x***_*n*_. The RNN module then made a final decision by accumulating *T*_*n*_ samples ({***k***_*n*_[0], ***k***_*n*_[1], …, ***k***_*n*_[*T*_*n*_]}) from ***e***_*n*_, where *T*_*n*_ (e.g., 8 or 16) simulated presentation time. A key aspect of SDNet is that the initial hidden state for processing ***x***_*n*_ (***h***_*n*_[0]) was inherited from the final hidden state of the *n −* 1th image (***h***_*n−*1_[*T*_*n−*1_]). The deterministic orientation was sampled from the RNN module’s final output evidence ***ϵ***_*n*_.

##### Behavioural data generation

For comparative analysis, 83 instances of SD-Net were created, each trained with different random splits of the image dataset. For each instances, four sequences of 50 test stimulus images (total 200 images per instance) were generated, mimicking the block structure of the human experiment. No instance-level screening was applied, as the instances were already designed to reproduce human-level precision. However, trial-level screening was performed, excluding samples with an orientation error exceeding 37.5^°^, consistent with the human data screening. This resulted in 15,717 samples (883 samples (approximately 5.3%) were excluded) from 83 SDNet instances for analysis.

#### 4.1.3 Analysis of serial dependence

##### Adjustment error and relative orientation

For each trial *n*, the stimulus properties (orientation *y*_*n*_, spatial frequency *f*_*n*_, phase *p*_*n*_, and presentation time *t*_*n*_) and the human participant’s /model instance’s response (*z*_*n*_) were recorded. The adjustment error *e*_*n*_ was calculated as the shortest angular distance between the stimulus and response orientation:

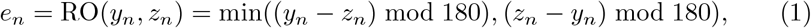

where min(· · ·) is an operator that outputs the minimum value of the input elements. The relative orientation from *m*-back trial, 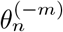, was defined as the shortest angular distance between the current stimulus orientation *y*_*n*_ and the stimulus orientation from *m* trials back *y*_*n−m*_:

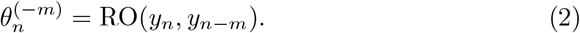

For each participant/instance, adjustment errors were averaged within bins defined by relative orientation *θ*, and other stimulus conditions (*f, t, f*_*−*1_, *t*_*−*1_ for spatial frequency, presentation time, and their 1-back counterparts):

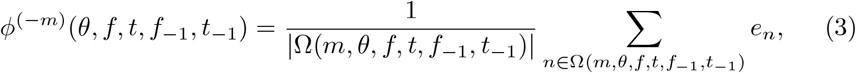

for *m* = 1, 2, 3. Here, Ω(*m, θ, f, t, f*_*−*1_, *t*_*−*1_) represents the set of trial indices satisfying the given conditions within a bin of width 10^°^ centered at *θ* given 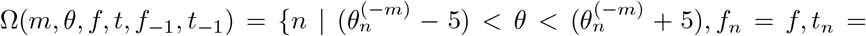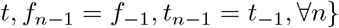.

##### Derivative of Gaussian (DoG) Fitting

To quantify the amplitude of serial dependence, the averaged error plot (*θ* versus *ϕ*^(*−m*)^(*θ, f, t, f*_*−*1_, *t*_*−*1_)) was fitted to a derivative of Gaussian (DoG) curve [20] for a given conditions {*m, f, t, f*_*−*1_, *t*_*−*1_}. The DoG function is defined as:

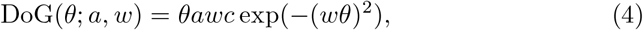

where *a* is the amplitude of the curve, representing the strength and direction of serial dependence, *w* is the width parameter describing the range of relative orientations over which the effect occurs, and 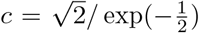 is a constant. Omitting the condition variables, *m, f, t, f*_*−*1_, *t*_*−*1_ from *ϕ*^(*−m*)^(*θ, f, t, f*_*−*1_, *t*_*−*1_) for simplicity notation, the parameters *a* and *w* were optimized by minimizing the squared error between the empirical adjustment errors *ϕ*(*θ*) and the DoG function across all participants/instances:

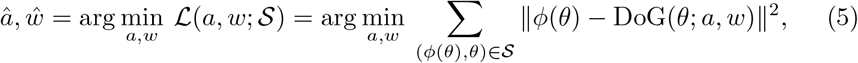

where 𝒮 is a set of pairs of a relative orientation error *θ* and corresponding adjustment error *ϕ*(*θ*) for all participants/instances. A sequential least squares programming method, implemented via `scipy.optimize.minimize` in Python, was used for this minimization.

##### SD amplitude quantification

Serial dependence was quantified by the SD amplitude (|SD|) and peak relative orientation (ΔSD) derived from the fitted DoG curve. The peak relative orientation is given by

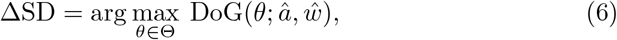

and the corresponding SD amplitude is

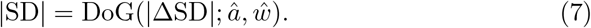

Here, Θ is the range of possible relative orientations, {−90.0, −89.9, −89.8, …, 89.8, 89.9} degrees. A positive amplitude indicates an attractive bias, while a negative amplitude would indicate a repulsive bias.

##### Statistical testing via bootstrap method

Statistical significance for the presence of serial dependence and for comparisons between conditions was assessed using a bootstrap method. To test for the presence of serial dependence in a given dataset 𝒮:

1. We generated 100,000 bootstrap samples 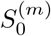, *m* = 1, …, 100000 by resampling with replacement from 𝒮. For each 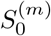, we performed DoG fitting to obtain loss 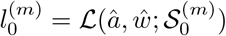.
2. To establish a null distribution, we randomly permuted the adjustment error values *ϕ*(*θ*) within each bin of 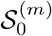 (while keeping *θ* fixed) to create 100,000 permuted sample set 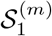. For each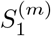, we obtained 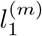.
3. The *p*-value for the null hypothesis (i.e., DoG function does not fit the data better than chance) was calculated as *p* = *K/*100000, where *K* is the umber of bootstrap samples where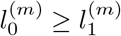.

To compare the strengths of serial dependence between two datasets, 𝒮_1_ and 𝒮_2_:

1. We generated 100,000 bootstrap samples 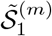 and 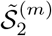 by resampling with replacement from 𝒮_1_ and 𝒮_2_, respectively.
2. DoG fitting was performed on each boostrapped sample to obtain 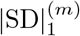 for 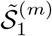 and 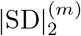 for 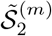.
3. For a one-sided test (e.g., whether 𝒮_1_ has stronger serial dependence than 𝒮_2_), the *p*-value was calculated as *p* = *K/*100000, where *K* is the count of samples where 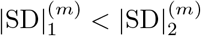.

### 4.2 Numerosity judgement task

In additon to the orientation adjustment task, we evaluated SDNet using a public dataset from a numerosity judgement task.

#### 4.2.1 Human behavioral dataset

We utilized a previously recorded dataset of 32 human participants who performed a numerosity judgement task [22]. In each trial, participants were sequentially presented with three images (inducer, reference, and probe) containing various numerosity of dots (8, 12, 16, 24, 32 dots), dot size (4, 6, 8 pixel), presentation durations (140, 200, 280 ms). Participants’ task was to determine whether the probe stimulus had more dots than the reference stimulus^1^. Each participant completed 10 blocks of 40 trials, totaling 400 trials. Taking breaks between the blocks, the participants completed 10 blocks, resulted in 400 trials done in total. Unlike the original study, no participants were excluded based on EEG signal quality for this behavioral analysis.

#### 4.2.2 SDNet

##### Architecture

The SDNet model for the numerosity judgement task (Figure 8) also consists of CNN and RNN modules. The CNN architecture was identical to that used for the orientation adjustment task (AlexNet-based). However, its output was a 37-dimensional probability distribution over dot numerosities (from 0 to 36). The RNN module was implemented with a GRU of *Q* = 8 hidden cells, accumulating samples from the CNN’s outout evidence.

**Figure 8.**
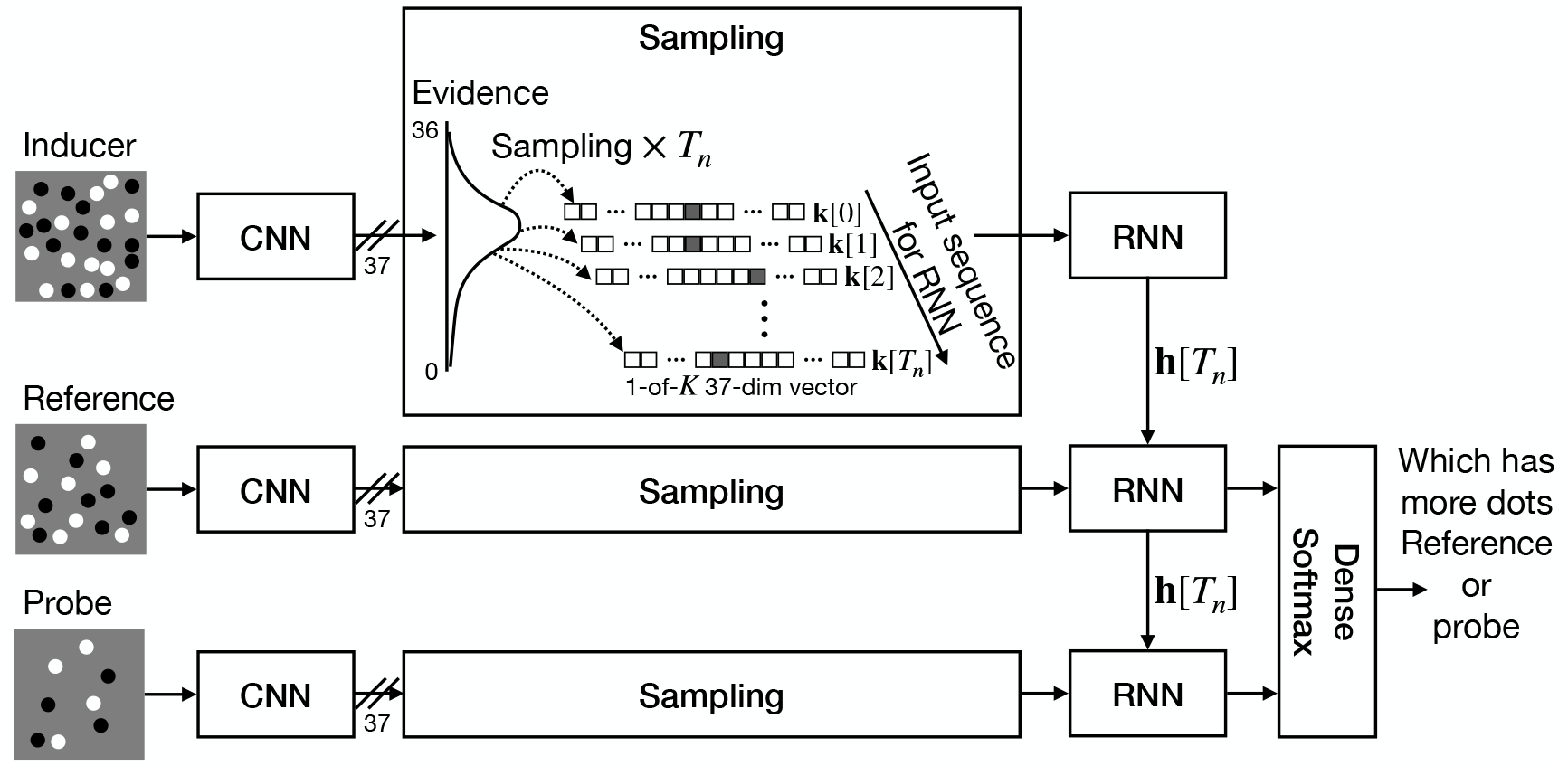
SDNet architecture for the numerosity judgement task. The model processes three sequential images (inducer, reference, probe) using a shared-weight CNN-RNN pathway. Each image is initially processed by a CNN module for feature extraction, and its evidence is accumulated by an RNN module. Crucially, the RNN’s hidden state for the reference image is initialized with the final hidden state from the inducer image, and similarly, the RNN’s hidden state for the probe image is initialized with the final hidden state from the reference image. The final hidden states from the reference and probe RNNs are then concatenated and fed into a judgement module (a dense layer followed by a softmax layer) that determines whether the probe image has more dots than the reference.

Since the task involved comparing two images (reference and probe), these were independently processed by separate CNN-RNN pathways. The final hidden states for the reference (***h***_ref_) and probe (***h***_probe_) images were concatenated to form a 16-dimensional feature vector (as *Q* = 8, so 8 × 2 = 16). This vector then fed into a judgement module, consisting of a single dense layer (reducing to 2 dimensions) and a softmax layer, to output the likelihood that the probe image had more dots.

##### Training

Training data consisted of 32,400 images of random dots, generated using the original script used from [22]. Dot numerosities ranged from 1 to 36, and dot sizes 2 to 10 pixels, covering the settings of the human experiment. Images were generated at 1080 × 1080 pixel resolution and then scaled to 128 × 128 for SDNet input, preserving dot size ratios. The dataset was split 6:4 into training (19,440 images) and validation (12,960 images) sets.

The CNN module was trained first to predict dot numerosity by minimizing cross-entropy loss. Early stopping was applied when the CNN’s average absolute numerosity error on the validation set reached below 3. This threshold was empirically determined to achieve human-level final judgement precision.

The RNN and judgement modules were then trained with the CNN weights fixed. The CNN’s output evidence ***e*** was sampled 16 times to generate sequences ***k***[*t*] (*t* = 0, …, 16) for RNN input. The RNN’s hidden state was updated as ***h***[*t*] = GRU(***k***[*t*], ***h***[*t −* 1]). For each training trial, inducer, reference, and probe images were randomly selected. Crucially, the initial hidden state for the reference image’s RNN was set to the final hidden state of the inducer image’s RNN, and the initial hidden state for the probe image’s RNN was set to the final hidden state of the reference image’s RNN. The RNN and judgement modules were trained to minimize the binary cross-entropy loss for the target variable *γ*, where *γ* = 1 if probe numerosity was greater than reference, and *γ* = 0 otherwise. Early stopping was applied when SDNet’s average judgement error (excluding equal numerosity trials) on the validation set reached below the human-averaged error (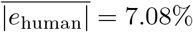)

##### Generating behavioural response

To generate SDNet behavioural data for comparison, 32 instances were trained using different dataset splits. For each trained instance, 400 trial sequences were generated using images from a separate test set, randomly combining inducer, reference, and probe images with parameters matching the human experiment (dot numerosity: 8, 12, 16, 24, 32; dot size: 4, 6, 8). This resulted in 400 responses per SDNet instance, matching the human dataset’s sample size.

#### 4.2.3 Analysis for serial dependence

##### Point of subjective equality (PSE)

For each participant/instance, behavioural data consisted of inducer (*a*_*n*_), reference (*b*_*n*_), probe (*c*_*n*_) dot numerosities, and the binary response *r*_*n*_ ∈ {0, 1} (0: reference had more dots, 1: probe had more dots) for trial *n*.

For specific inducer and reference numerosities (reference was fixed at 16 dots in this dataset), the proportion of “probe has more dots” responses, *ψ*(*c*), was calculated for each probe numeoristy *c*:

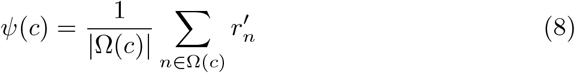

where Ω(*c*) = {*n* | *c*_*n*_ = *c, ∀n*} defines the set of trials for a given probe numerosity.

A psychometric function (Weibull function) *F* (*θ*; *α, β*) was fitted to these proportions across all participants/instances by minimizing the squared error:

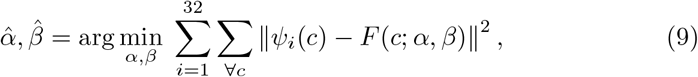

where *ψ*_*i*_(*c*) is the proportion for the *i*th participant/instance.

PSE was defined as the probe numerosity *c* at which the psychometric function yielded a 0.5 probability of responding “probe has more dots”:

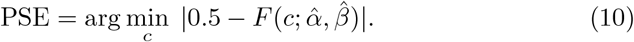

Serial dependence strength was quantified by analyzing shifts in PSE across different inducer numerosities.

##### Statistical testing via bootstrap method

Similar to the orientation adjustment task, a bootstrap method was employed for statistical testing of PSE differences. To compare PSEs between two conditions (e.g., different inducer numerosities for 𝒮_1_ and 𝒮_2_):

1. We generated 100,000 bootstrap samples 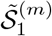 and 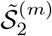 by resampling with replacement from 𝒮_1_ and 𝒮_2_, respectively.
2. For each bootstrapped sample, PSEs were computed as 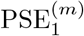 from 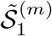 and 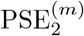from 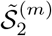.
3. For a one-sided test, the *p*-value for testing whether PSEs from 𝒮_1_ and 𝒮_2_ are significantly different was calculated as *p* = *K/*100000, where *K* is the count of samples where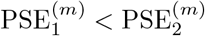.

## Acknowledgements

This work was supported in part by the Japan Society for the Promotion of Science (JSPS) KAKENHI, grant number 22H05163 and 24K15047, and Japan Science and Technology Agency (JST) Advanced International Collaborative Research Program (AdCORP), grant number JPMJKB2307.

## Author contributions

H.H. and K.H. designed and performed the research, collected and analyzed the data. H.H drafted the paper. H.H, K.H., and Y.T edited the paper.

## Competing interests

The authors declare no competing interests.

## Footnotes

1 The experiments also included two another tasks. One of the participants’ tasks was to determine whether the probe stimulus had bigger dots than the reference stimulus. The other was to determine whether the probe stimulus had longer presentation time than the reference stimulus. In our experiment, we did not use these data.

## Notes

### Competing Interest Statement

The authors have declared no competing interest.

